# Effects of environmental manipulations on cocaine-vs-social choice in male and female rats

**DOI:** 10.1101/2022.07.15.500221

**Authors:** Madison M. Marcus, S. Stevens Negus, Matthew L. Banks

**Affiliations:** Department of Pharmacology and Toxicology, Virginia Commonwealth University School of Medicine, Richmond, VA, USA 23298

**Keywords:** cocaine, social interaction, choice, progressive ratio, addiction, self-administration

## Abstract

Cocaine use disorder occurs in an environment where cocaine and other nondrug commodities are concurrently available. Preclinical drug-vs-nondrug choice procedures are one simplified method of modeling this complex clinical environment. The present study established a discrete-trial progressive-ratio (PR) cocaine-vs-social interaction choice procedure in male and female rats and determined sensitivity of choice behavior to manipulations of reinforcer magnitude and non-contingent “sample” reinforcer presentation. Rats could make up to nine discrete choices between an intravenous cocaine infusion (0.1 – 1.0 mg/kg/inf) and social interaction with a same-sex social “Partner” rat. Cocaine infusions were available under a PR schedule of reinforcement, and social interaction was available under a fixed-ratio (FR) 3 schedule. Social interaction was chosen over no or small cocaine doses (saline, 0.01 mg/kg/inf) and behavior was reallocated away from social and towards cocaine at larger cocaine doses (1.0 mg/kg/inf). Manipulating social interaction time as one method to alter social reinforcer magnitude did not significantly alter cocaine-vs-social choice. Removing the non-contingent reinforcer presentations before the discrete choice trials also failed to affect cocaine-vs-social choice, suggesting the time interval was sufficient to minimize any potential influence of the non-contingent cocaine infusions on subsequent choice behavior. Overall, the present results were consistent with previous drug-vs-social choice studies and extend our knowledge of environmental factors impacting drug-vs-social choice. Future studies determining the pharmacological sensitivity of cocaine-vs-social choice will be important in expanding the preclinical utility of these procedures for candidate medication drug development.

## 1.0 Introduction

Cocaine use disorder is a global public health problem. The 2019 National Survey on Drug Use and Health reported that over 1 million people met criteria for cocaine use disorder, and cocaine-related overdose deaths have been increasing every year since 2010 (Jones et al., 2017; Lipari and Park-Lee, 2020). Currently, there are no Food and Drug (FDA)-approved pharmacotherapies to treat cocaine use disorder. Preclinical research is one pathway towards improving our fundamental knowledge of the mechanisms associated with cocaine abuse and facilitating evaluation of candidate treatment strategies.

Substance use disorders are increasingly being recognized as a maladaptive allocation of behavior between drug use and other behaviors maintained by more adaptive non-drug reinforcers such as social interaction, food, or money. Preclinical drug-choice self-administration procedures are one category of behavioral procedures that have improved our understanding of behavioral allocation or “choice” between addictive drugs and nondrug reinforcers. Decades of preclinical research have demonstrated that cocaine-vs-food choice is sensitive to environmental manipulations such as reinforcer magnitude (Banks and Negus, 2012; Cantin et al., 2010; Nader and Woolverton, 1991; Negus, 2003; Thomsen et al., 2013), cost (Cantin et al., 2010; Nader and Woolverton, 1992; Negus, 2003; Thomsen et al., 2013), and punishment (Negus, 2005) in both rats and nonhuman primates. These preclinical results are congruent with human laboratory cocaine-vs-money choice studies (Foltin et al., 2015; Hart et al., 2000; Lile et al., 2016).

One non-drug reinforcer readily available in the clinical environment is social interaction (e.g., family and friends). For example, one effective therapeutic strategy in treating substance use disorders is the “community reinforcement approach,” in which drug abstinence is reinforced through increased interaction with family and community members (Meyers et al., 2011). Recently, social interaction was established as a reinforcer in a preclinical drug-vs-social “choice” procedure in rats (Venniro et al., 2018), extending previous nonhuman primate research on food-vs-social choice (Corwin and Schuster, 1993). Results from these drug-vs-social choice studies demonstrated that social interaction as a concurrently available nondrug reinforcer produced methamphetamine, cocaine, and heroin abstinence (Venniro et al., 2021, 2018); however, choice behavior was reallocated away from social and towards cocaine following both delay of social reinforcement and increases in social-reinforcer price (Venniro et al., 2021). The sensitivity of cocaine-social choice to reinforcement delay and reinforcer price supports the further investigation of additional environmental variables influencing cocaine-vs-social choice.

The goal of the current study was to establish a discrete-trial cocaine-vs-social choice procedure in rats modeled after previously established cocaine-vs-food choice procedures in monkeys and humans (Johnson et al., 2016; Lile et al., 2016). Once established, two reciprocal environmental variables were manipulated: (1) cocaine dose, to evaluate behavioral sensitivity to cocaine-reinforcer magnitude, and (2) social-access duration, as a strategy to manipulate social-reinforcer magnitude. In addition, drug self-administration choice procedures sometimes include non-contingent reinforcer presentations during a “sample” period before the subsequent choice components. These initial reinforcer presentations are intended to serve as part of a compound discriminative stimulus that also includes non-drug cues (e.g., stimulus lights) for subsequent choice behavior (Thomsen et al., 2013; Townsend et al., 2021); however, non-contingent drug injections may also produce other effects that bias responding toward drug choice and exaggerate metrics of drug preference (Vandaele et al., 2016). Accordingly, a final goal of this study was to compare the effects of including or omitting the presentation of non-contingent cocaine and social stimuli at the start of each behavioral session.

## 2.0 Material and Methods

### 2.1 Subjects

A total of 16 Sprague-Dawley rats (8 males and 8 females) weighing 230-300g upon arrival were acquired (Envigo, Frederick, MD). Eight rats (4 male and 4 female) were designated as “Responder” rats and paired with a same sex social “Partner” rat. Final sample sizes for each experiment ranged from five to seven and are reported in figure legends. Responder rats were pair housed with their designated social partner for one week upon arrival before being separated into individual housing for training and testing during the remainder of the study. Animals were housed in a temperature and humidity-controlled vivarium that was maintained on a 12-h light/dark cycle (lights off at 6:00 PM). Water and food (Teklad Rat Diet, Envigo) were provided ad-libitum in the home cage throughout all experiments. Behavioral sessions were conducted five days per week from approximately 11am – 3pm. Animal maintenance and research were conducted in accordance with the 2011 Guidelines of the National Institutes of Health Committee on Laboratory Animal Resources, and both enrichment and research protocols were approved by the Virginia Commonwealth University Institutional Animal Care and Use Committee.

### 2.2 Apparatus and Catheter Maintenance

Modular drug-social operant chambers (Med Associates, St. Albans, VT) were based on a published design (Venniro and Shaham, 2020). Briefly, modular operant chambers were located within sound-attenuating cubicles and equipped with two retractable levers on opposite chamber walls. Mounted above each lever was a white stimulus light that served as a visual discriminative stimulus for availability of each reinforcer. Response requirement completion on the cocaine-paired lever resulted in intravenous (IV) cocaine delivery by a syringe pump (PHM-100, Med Associates) located inside the sound-attenuating cubicle. The social “Partner” rat was placed in an adjoining chamber, separated from the “Responder” rat by a sliding guillotine door. Response requirement completion on the social-paired lever resulted in the retraction of the guillotine door, allowing for access to the “Partner” rat behind a metal barrier. As previously described, the barrier contained cut-out sections that allowed visual and tactile contact between the two rats, but prevented comingling. Opening and closing the guillotine door allowed for precise control of the duration of the social-access period and automation of behavioral procedures (Venniro and Shaham, 2020). Rats were surgically implanted with a custom-made jugular catheter and vascular access port as previously described in detail (Townsend et al., 2021). Rats were given five recovery days before beginning behavioral testing. After each behavioral session, catheters were flushed with 0.1 ml gentamicin (0.4 mg/ml) followed by 0.1 ml of heparinized saline (30 U/ml). Catheter patency was verified periodically and at the end of each experiment by muscle tone loss within 5 s of IV methohexital (0.5 mg) administration.

### 2.3 Cocaine and Social Choice Training

#### 2.3.1 Single operant training

Rats were first trained to respond for 30-s social interaction under a fixed-ratio (FR) 1 schedule of reinforcement during daily 30-min behavioral sessions. Each session began with illumination of the house light, extension of the social-paired lever, and illumination of the social-paired stimulus light. Following each response-requirement completion, the lever retracted, and the guillotine door opened for 30 s. The stimulus and house lights remained illuminated in the chamber throughout the procedure. This schedule of reinforcement was maintained until the “Responder” rat earned ≥ 10 reinforcers per 30-min session for at least three consecutive days. Once acquisition criteria were met, the FR requirement was increased over consecutive days until the terminal FR3 schedule was reached. If a rat failed to respond within six days, the “Responder” and “Partner” rat positions were switched such that the previously assigned “Partner” rat was now designated the “Responder” rat. Rats remained on the FR3 schedule until at least five reinforcers were earned during the 30-min session. After meeting social self-administration acquisition criteria, rats were implanted with IV jugular catheters and trained to respond for IV cocaine infusions (0.32 mg/kg/inf) under an initial FR1 / 15-s time out schedule of reinforcement during daily 2-h behavioral sessions. Each session began with a non-contingent cocaine infusion followed by a 60-s time out. The response period was signaled by extension of the drug-associated lever and illumination of the drug-paired stimulus light above the lever. Following each response-requirement completion, the lever retracted, the stimulus light was extinguished, and an IV drug infusion was administered. Once rats self-administered ≥ 30 cocaine infusions during a 2-hr session, the FR requirement was increased to FR3. Rats remained on the FR3 schedule until ≥ 30 drug infusions were earned during the session.

#### 2.3.2 Cocaine-vs-Social Choice Training

Once both social- and cocaine-maintained responding had been established separately, choice behavior was established in rats using a discrete-trial cocaine-vs-social choice procedure. The purpose of this training phase was to acclimate rats to discrete-trial choice procedures and cocaine dose manipulations. During training, both social and cocaine reinforcers were available under FR3 schedules of reinforcement. Each session began with a 12-min “sample” component, followed by five 10-min discrete-choice trials. The sample component began with illumination of the social cue light and retraction of the guillotine door to allow 30 s of social interaction with the “Partner” rat. After 30 s, the guillotine door lowered, all stimulus lights extinguished, and a 5-min time out period began. Next, the cocaine cue light was illuminated, and the rat was administered a non-contingent infusion of the unit cocaine dose available during the subsequent choice trials. After the cocaine infusion, all stimulus lights were extinguished, and a 5-min time out started. During each subsequent discrete-choice trial, both levers were extended, and stimulus lights above both levers were illuminated to signal concurrent availability of both the social and cocaine reinforcers. The first lever press emitted during each choice trial locked in the rat’s choice for that given trial, deactivated, but did not retract, the alternate lever, and extinguished the alternate stimulus light. Responses on the deactivated lever were recorded but had no programmed consequences. Ratio-requirement completion on the active lever before the 10-min trial had elapsed resulted in delivery of the associated consequent stimulus, retraction of both levers, and offset of both stimulus lights for the remainder of that 10-min choice trial. If the rat failed to meet the response requirement on the active lever before 10 min elapsed, both levers retracted, both stimulus and house lights were extinguished, and the trial was counted as an omission. Each of the five trials was separated by a 30-s time out period. Response-requirement (FR3) completion on the social-paired lever resulted in 30-s access to the “Partner” rat, whereas response requirement (FR3) completion on the cocaine-associated lever resulted in delivery of the unit cocaine dose available for that session. Cocaine dose (0.1, 0.32, 1.0 mg/kg/inf) was varied across sessions by changing the infusion duration (e.g., 300g rat; 1.5, 5, 15-s of pump activation, respectively), and cocaine doses and saline were presented in a random order. Each cocaine dose was evaluated for at least three and up to five consecutive sessions until choice behavior was considered stable as defined by either (a) the number of trials completed for each reinforcer fluctuating by no more than one trial for two consecutive days, or (b) the rat showing preference for one reinforcer for three or more consecutive days. Data from the final two training days were averaged for both graphical and statistical purposes (see Supplemental Materials).

### 2.4 Progressive-Ratio Cocaine-vs-Social Choice

Following training completion, rats transitioned to the terminal progressive-ratio (PR) discrete-trial choice procedure. This procedure was modeled after aspects of a discrete-trial drug-vs-food choice procedure used in both monkeys and humans (Johnson et al., 2016; Lile et al., 2016). Specifically, daily sessions began with the sampling component as described in section 2.3.2, and then proceeded to nine 10-min choice trials. The response requirement for cocaine increased in quarter-log increments (FR 3, 6, 10, 18, 32, 56, 100, 180, 320) after each completed cocaine choice. The procedure consisted of nine choice trials to permit high response requirements that captured the cocaine breakpoint. Since prior literature indicates that social interaction maintains lower levels of operant responding relative to food reinforcers (Baldwin et al., 2022; Chow et al., 2022), the ratio requirement on the social lever was held constant at FR3 during all trials to facilitate maintenance of social responding. If a rat failed to complete a ratio requirement within the 10-min trial period, the trial terminated, a 30-s time out period ensued, the trial was counted as an omission, and the ratio requirement did not increment for the next trial.

Using the terminal progressive-ratio cocaine-vs-social choice procedure, three variables were manipulated: cocaine dose, social-access time, and presence/absence of non-contingent reinforcer presentation. In the first experiment, saline and three cocaine doses (0.1, 0.32, 1.0 mg/kg/inf) were tested to determine a cocaine choice dose-effect function. Cocaine dose was varied by changing the infusion duration as described in section 2.3.2, and cocaine doses and saline were evaluated in a random order. A second experiment attempted to manipulate the magnitude of the social reinforcer by varying the duration of the social interaction period. In this experiment, the cocaine dose was 1.0 mg/kg/inf as the alternative to 60-s social interaction, or 0.32 mg/kg/inf as the alternative to 0-s social interaction. In the latter condition, the “Partner” rat was not placed in the adjoining chamber to eliminate any auditory or olfactory stimuli associated with “Partner” rat presence, and response requirement completion on the social-lever resulted in retraction of both levers and offset of all stimulus lights but did not open the guillotine door. A third and final experiment determined whether non-contingent reinforcer presentations during the sampling period were determinants of subsequent choice behavior, as suggested by the literature using non-discrete-trial choice procedures (Thomsen et al., 2013; Townsend et al., 2021). The non-contingent reinforcer presentations (i.e., cocaine infusion and social interaction) were removed from the sampling component, leaving only the illumination of stimulus lights to signal reinforcer availability. The rest of the procedure remained the same as previously described. In this experiment, the cocaine dose was 1.0 mg/kg/inf cocaine as the alternative to 30-s social interaction during each of the nine discrete-trial choice components. For all three experiments, independent-variable manipulations were examined for four to five consecutive days, and the data from the final two days were averaged within a rat and then across rats for both graphical and statistical purposes.

### 2.5 Data Analysis

The primary dependent measures in the discrete-trial cocaine-vs-social choice procedure were (1) number of trials completed for each reinforcer and omitted trials, and (2) percent cocaine choice, defined as [(number of ratio requirements, or “choices,” completed on the cocaine-associated lever / total number of choices completed on both the cocaine- and social-associated levers) x 100]. These measures were plotted as a function of cocaine dose or independent-variable manipulation. Data were analyzed using repeated-measures one-way or two-way analysis of variance, or mixed-effects analysis as appropriate. Sphericity violations were corrected using the Geisser-Greenhouse epsilon. Significant main effects or interactions were followed by planned post-hoc tests that corrected for multiple comparisons. The criterion for significance was set a priori at the 95% level of confidence (p<0.05), and all analyses were conducted using GraphPad Prism (v 9.2.0, La Jolla, CA).

### 2.6 Drugs

(-)-Cocaine HCl was provided by the National Institute on Drug Abuse Drug Supply Program (Bethesda, MD, USA). Cocaine was dissolved in bacteriostatic sterile saline and passed through a 0.22-micron sterile filter before IV administration. Cocaine doses are expressed as the salt form listed above.

## 3.0 Results

### 3.1 Cocaine and Social Choice Training

Data on training phases are shown in Supplemental Table 1 and Figures S1-5. Seven rats completed all required training and proceeded to experiments under the terminal PR cocaine-vs-social choice procedure.

### 3.2 PR Cocaine-vs-social choice

Figure 1 shows the cocaine-vs-social choice dose-response function when the cocaine response requirement increased across trials according to a PR schedule and the social response requirement was FR3. Figure 1A shows cocaine choice was sensitive to cocaine dose, such that 1 mg/kg/inf cocaine maintained greater percent cocaine choice compared to saline (Dose: F(1.6, 7.1) = 14.2, p = 0.0041). No sex differences were detected, and cocaine choice data separated by sex are shown in Supplemental Figure S6. Figure 1B shows the number of cocaine and social trials completed and trials omitted varied as a function of cocaine dose (Reinforcer: F(1.7, 10.4) = 58.9, p = <0.0001; Dose × Reinforcer: F(2.3, 8.0) = 17.3, p = 0.001). When saline (0 cocaine dose) or 0.1 mg/kg/inf cocaine was available, rats earned less than three infusions and completed significantly more trials on the social lever than the cocaine-associated lever. Conversely, at the 1.0 mg/kg/inf cocaine dose, rats earned nearly six cocaine infusions and completed significantly more trials on the cocaine-lever than the social-lever. Omissions were low across all cocaine doses. Figure 1C-D show individual subject data for each of the nine discrete trials during the last behavioral session where 0.1 mg/kg/inf (Fig. 1C) or 1.0 mg/kg/inf (Fig. 1D) cocaine was available. At both cocaine doses, all but one rat (subject F4 during 1.0 mg/kg/inf cocaine availability) switched between responding on the cocaine- and social interaction-paired levers, sampling both reinforcers throughout the session.

**Figure 1.**
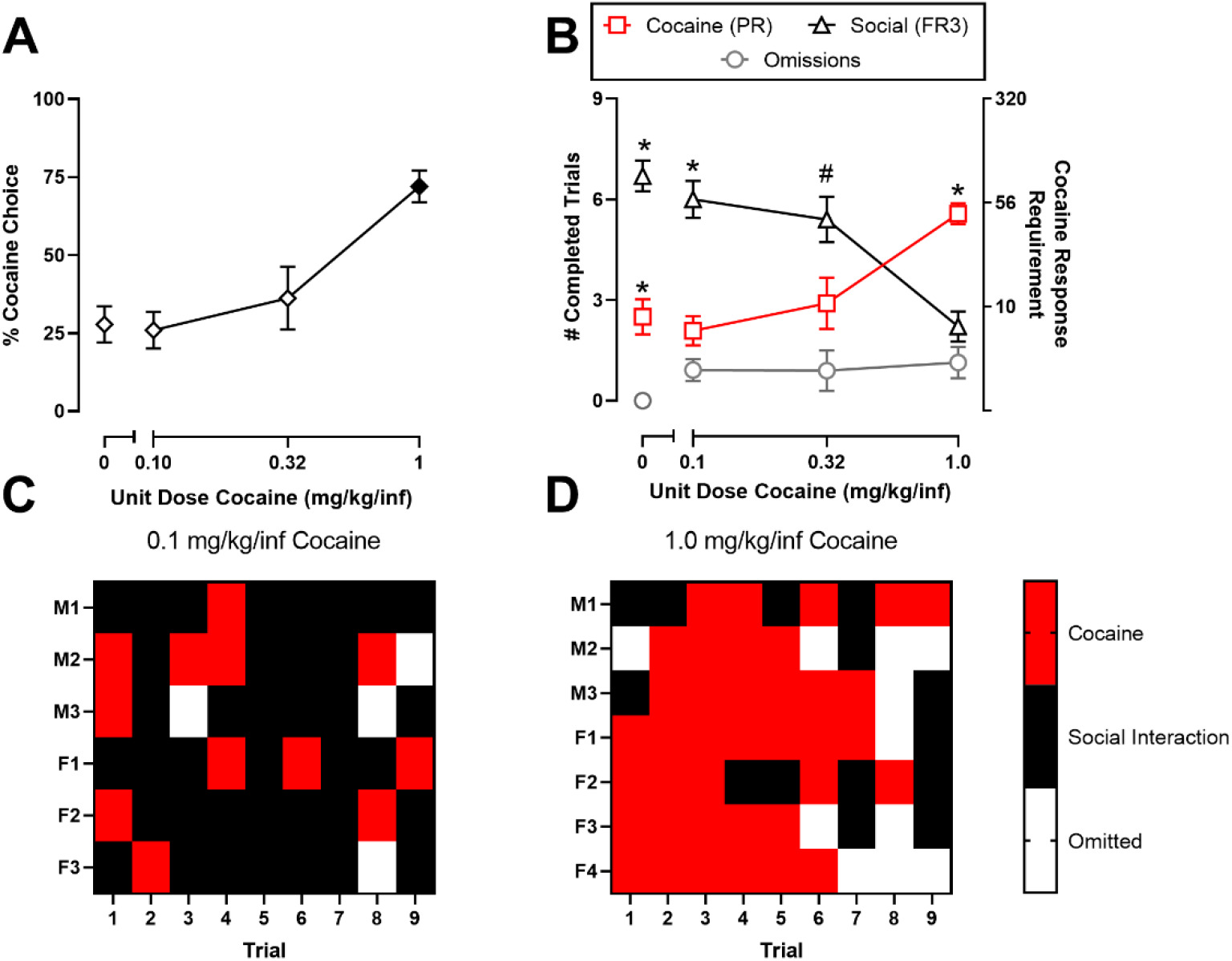
Effect of cocaine dose on cocaine-vs.-social interaction choice. Top panels: (A) percent cocaine choice as a function of cocaine dose. (B) trials completed for cocaine (PR 3, 6, 10, 18, 32, 56, 100, 180, 320), completed for social interaction (FR3), or omitted as a function of cocaine dose. All points represent the mean ± SEM for the final 2 days of testing. Filled points denote statistical significance (p < 0.05) relative to saline; symbols represent significant (p < 0.05) comparisons within a dose: *difference from the other reinforcer and omitted trials; ^#^difference from omitted trials only. Bottom panels: Individual subject behavioral allocation on the final day of testing between cocaine (red), social interaction (black), or omitted trials (white) when the alternative to social interaction was 0.1 mg/kg/inf cocaine (C) or 1.0 mg/kg/inf cocaine (D). Each row corresponds to a single subject, and each column corresponds to a discrete choice trial. Results depicted are representative of one testing day. n = 5 (2M/3F) for saline, n = 6 (2M/4F) for 0.10 mg/kg/inf cocaine, n = 5 (2M/3F) for 0.32 mg/kg/inf, and n = 7 (3M/4F) for 1.0 mg/kg/inf cocaine.

Figure 2 shows the time course of cocaine choices, social choices, and omissions per session across days of availability at each cocaine dose. Social interaction was chosen over saline across all five testing days (Fig. 2A; Reinforcer: F(1.0, 4.1) = 36.6, p = 0.0035). When 0.1 mg/kg/inf cocaine was available as the alternative to social interaction, rats completed more social than cocaine trials only on day 5 (Fig. 2B; Reinforcer: F(1.9, 9.7) = 10.5, p = 0.004; Time × Reinforcer: F(2.1, 7.2) = 5.4, p = 0.036). Trials completed during 0.32 mg/kg/inf cocaine and social availability was stable across the 5 days with no significant difference between the two reinforcers (Fig. 2C; Reinforcer: F(1.9, 7.8) = 45.26, p = 0.03). When 1.0 mg/kg/inf cocaine was available, the number of cocaine trials completed was significantly higher than the number of social trials completed on days 4 and 5 (Fig 2D; Reinforcer: F(1.4, 8.4) = 17.6, p = 0.0017; Time × Reinforcer: F(3.1, 17.2) = 3.9, p = 0.026). Across all experimental conditions, the number of omitted trials remained low.

**Figure 2.**
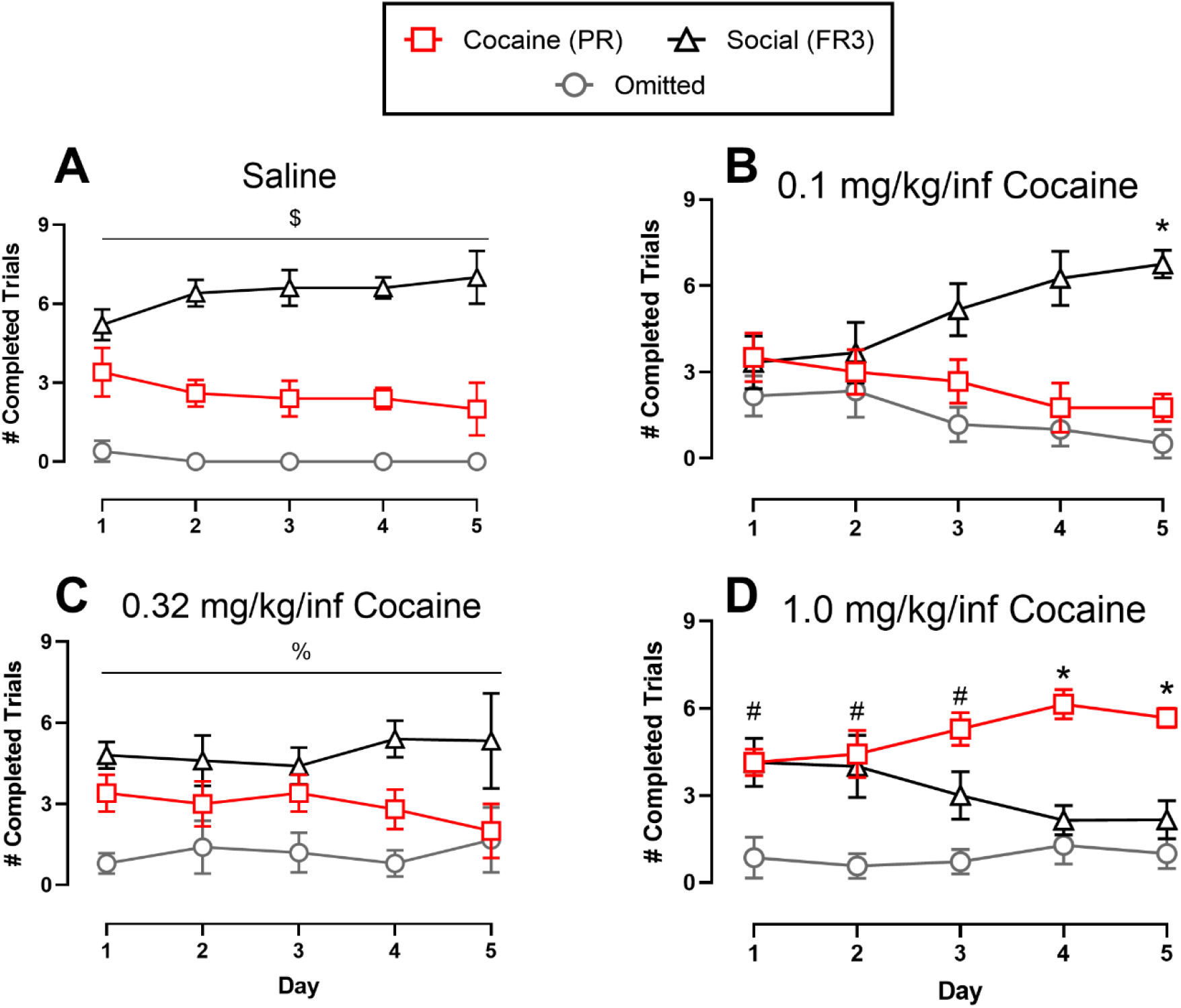
Cocaine-vs-social interaction choice across testing days. Abscissae: testing day. Ordinates: number of trials completed for cocaine, completed for social interaction, or omitted within a session. All points and bars represent the mean ± SEM. Lines represent main effect of Reinforcer (p < 0.05); Symbols represent significant (p < 0.05) post-hoc comparisons: $difference between cocaine, social, and omitted; *difference from the other reinforcer and omitted; %difference between social and omitted; ^#^Difference from omitted trials only. n = 2-5 (2M/3F) for saline, n = 4-6 (2M/4F) for 0.1 mg/kg/inf cocaine, n = 3-5 (2M/3F) for 0.32 mg/kg/inf cocaine, and n = 6-7 (3M/4F) for 1.0 mg/kg/inf cocaine dose.

### 3.3 Effect of manipulating social interaction access time on cocaine choice

After completion of the dose-effect curves (Section 3.2), behavioral allocation between 1.0 mg/kg/inf cocaine and 30-s social interaction was redetermined for five consecutive days. Percent cocaine choice from the final 2 days of testing is shown in Figure 3A (time course data not shown). Although percent cocaine choice appeared to have decreased from initial determination (Fig. 1A), this difference was not statistically significant. Increasing social interaction time to 60-s failed to significantly decrease cocaine choice compared to 30-s social interaction (Fig. 3A). Figure 3B shows the number of trials completed for both reinforcers remained stable across testing days (Reinforcer: F(1.0, 4.1) = 12.6, p = 0.023), with rats completing more trials for both reinforcers than omitting trials. However, there was no significant difference between the number of trials completed for cocaine and social interaction. Figures 3C-D show individual subject data for the last behavioral session when social interaction was 30-s (Fig. 3C) or 60-s (Fig. 3D). There were no consistent trends or patterns of behavior with all subjects completing both cocaine and social trials during the session.

**Figure 3.**
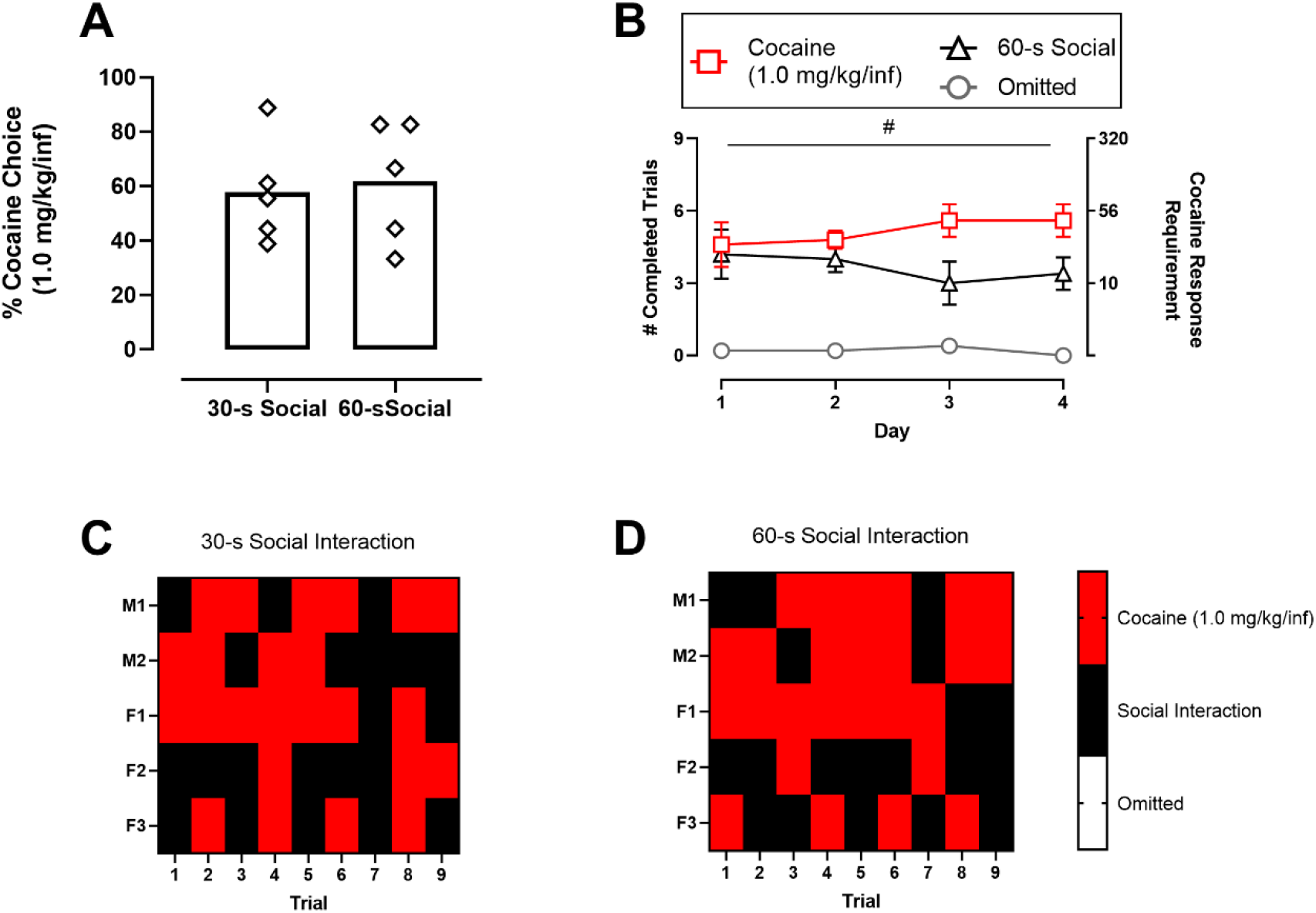
Effect of increasing social interaction duration on cocaine-vs-social choice. (A) Percent cocaine choice between 1.0 mg/kg/inf cocaine and 30-s or 60-s social interaction. Points represent individual subject means for the final 2 days of testing; bars represent group means. (B) number of trials completed for each reinforcer or omitted across testing days. Points and bars represent the daily mean ± SEM. Lines represent main effect of Reinforcer; ^#^difference from omitted trials only (p < 0.05) (C, D) Individual subject behavioral allocation on the final day of testing between cocaine (red), social interaction (black), or omitted trials (white) when the alternative to cocaine was 30-s (C) or 60-s social interaction (D)

Figure 4 shows that decreasing social interaction time from 30 s to 0 s by removing the “Partner” rat and keeping the guillotine door closed did not significantly increase cocaine choice compared to 30-s social interaction (Fig. 4A). There was no significant difference between the number of trials completed for cocaine, 0-s social interaction, or omitted across testing days (Fig. 4B). Figures 4C-D show individual subject data for the last behavioral session when social interaction was 30 s (Fig. 4C) or 0 s (Fig. 4D), highlighting individual subject variability. For example, subject F1 displayed exclusive social choice when social-interaction time was 30 s and failed to complete any trials when social-interaction time was reduced to 0 s. Conversely, both male subjects M1 and M2, and female subject F2 responded for both cocaine and social interaction when social-interaction time was 30 s and increased the number of cocaine trials completed when social-interaction time was decreased to 0 s.

**Figure 4.**
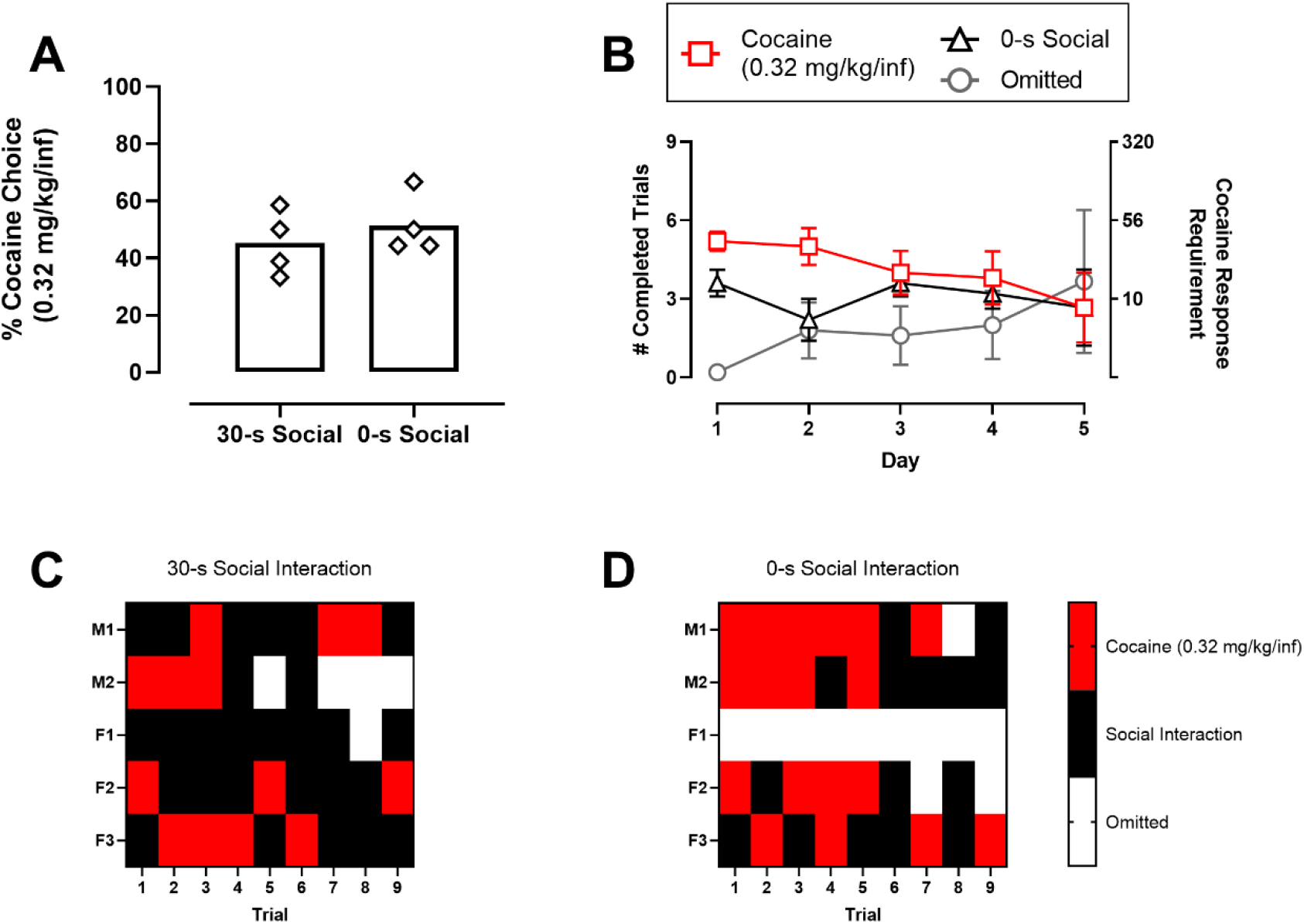
Effect of removing social interaction on cocaine-vs-social choice. (A) Percent cocaine choice between 0.32 mg/kg/inf cocaine and 30-s or 0-s social interaction. Points represent individual subject means for the final 2 days of testing; bars represent group means. (B) number of trials completed for each reinforcer or omitted across testing days. Points and bars represent the daily mean ± SEM. (C, D) Individual subject behavioral allocation on the final day of testing between cocaine (red), social interaction (black), or omitted trials (white) when the alternative to cocaine is 30-s (C) or 0-s social interaction (D)

### 3.4 Effect of non-contingent reinforcer presentation on cocaine choice

Figure 5 shows that eliminating non-contingent reinforcer presentations during the “sampling” period had no effect on choice between 1.0 mg/kg/inf cocaine and 30 s of social interaction. Percent cocaine choice was not affected by removal of non-contingent reinforcer presentations (Fig. 5A). Across testing days, more trials were completed for both reinforcers than omitted (Fig. 5B Reinforcer: F(1.1, 4.4) = 16.3, p = 0.01). Figures 5C-D show no robust changes in the behavior of individual subjects when the reinforcers were presented during the sampling period (Fig. 5C) compared to when they were absent (Fig. 5D). Consistent with prior testing conditions, (Fig. 3 & 4 C-D), individual rats alternated between responding for cocaine and social reinforcers across trials.

**Figure 5.**
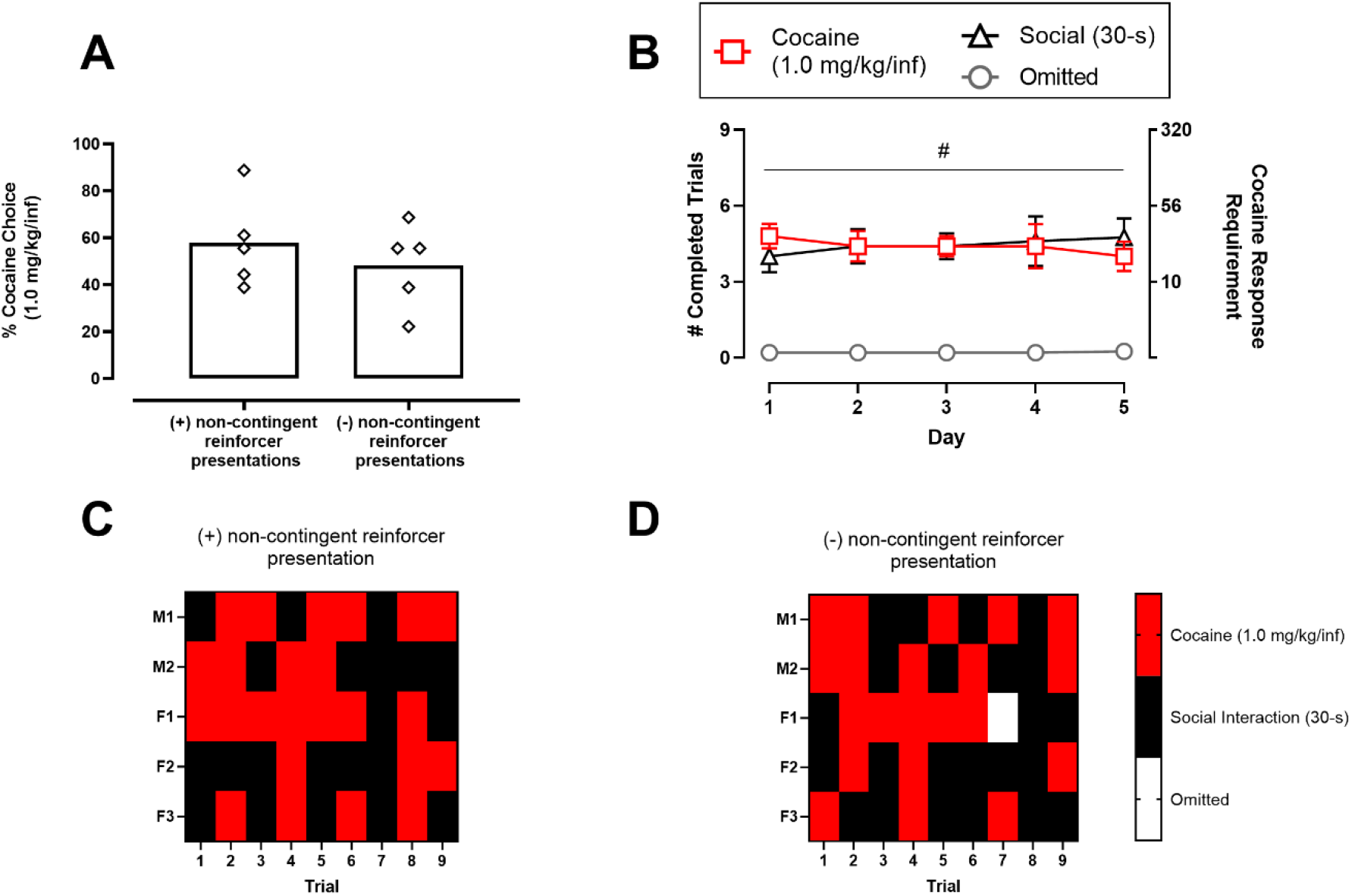
Effect of eliminating non-contingent reinforcer presentation on cocaine-vs-social choice. (A) Percent cocaine choice between 1.0 mg/kg/inf cocaine and 30 s social interaction with and without non-contingent cocaine and social presentation during the pre-choice sampling period of each daily session. Points represent individual subject means for the final 2 days of testing; bars represent group means. (B) number of trials completed for each reinforcer or omitted across testing days. Points and bars represent the daily mean ± SEM. Bars represent main effect of Reinforcer; ^#^difference from omitted trials only (p < 0.05) (C, D) Individual subject behavioral allocation on the final day of testing between cocaine (red), social interaction (black), or omitted trials (white) with (C) or without non-contingent reinforcer presentation (D)

## 4.0 Discussion

The present study established a progressive ratio discrete-trial cocaine-vs-social choice procedure in male and female rats. The sensitivity of this procedure to manipulations of reinforcer magnitude and non-contingent reinforcer presentation was determined. There were three main findings. First, increasing unit cocaine dose resulted in greater behavioral allocation towards cocaine and away from social interaction when cocaine was available under a PR schedule and social reinforcement was under an FR3 schedule. These results were in accordance with previous cocaine-vs-food choice studies in rats, nonhuman primates, and humans, where large cocaine doses are chosen over concurrently available non-drug reinforcers (Hart et al., 2000; Iglauer et al., 1976; Johnson et al., 2016; Lile et al., 2016; Thomsen et al., 2013). Second, in contrast to cocaine dose manipulations, experimental manipulations intended to alter the magnitude of social interaction as a reinforcer did not significantly alter cocaine-vs-social choice. These results were concordant with recent results demonstrating that social-access time was not a significant factor in operant responding for social interaction (Baldwin et al., 2022; Chow et al., 2022). Finally, non-contingent “sample” presentation of reinforcers before the discrete-choice trials did not significantly impact cocaine-vs-social choice. Furthermore, examination of individual subject data indicates that the majority of rats alternated choosing between the two concurrently available reinforcers within a behavioral session. These individual results do not support the hypothesis that cocaine intoxication is a major determinant of choice behavior. Overall, these results are generally consistent with and extend previous studies of social interaction as a reinforcer in rats. These results also support the utility of cocaine-vs-social choice to improve our basic understanding of behavioral allocation between cocaine and nondrug reinforcers.

### 4.1 Cocaine was chosen over social interaction under a PR schedule

Social interaction was chosen over no or small cocaine doses, and a large cocaine dose was chosen over social interaction. The sensitivity of cocaine-vs-social interaction choice to manipulations of drug reinforcer magnitude is consistent with previous cocaine-vs-food choice results using discrete-trial choice procedures (Johnson et al., 2016; Lile et al., 2016; Nader and Woolverton, 1992), non-discrete-trial choice procedures (Negus, 2003; Thomsen et al., 2013), and opioid-vs-social interaction choice studies (Chow et al., 2022).

### 4.2 Manipulation of social access time failed to alter cocaine-vs-social choice

Altering the magnitude of food as the non-drug alternative reinforcer in cocaine-vs-food choice procedures has reliably affected behavioral allocation between the drug and food reinforcer (Nader and Woolverton, 1991; Negus, 2003; Thomsen et al., 2013; Townsend et al., 2021). Therefore, we hypothesized that increasing social-reinforcer magnitude by increasing the social-interaction time would increase social choice and decrease cocaine choice. However, there were no significant differences in behavioral allocation when social-interaction time was increased from 30 to 60 s. The present results are consistent with other recent results also showing that increasing social-interaction time from 30 to 60 s was ineffective to increase social-reinforcer magnitude (Baldwin et al., 2022; Chow et al., 2022). Results of prior studies suggest the opportunity for social “rough and tumble” play is more reinforcing than generalized social interaction, as evidenced by rats only showing conditioned place preference for a “partner” rat that can participate in play behavior (Calcagnetti and Schechter, 1992; Trezza et al., 2009). In the current study, rats were separated by a metal barrier during the social interaction period. This allowed for automation of the choice procedure but prevented any opportunity for “rough and tumble” play between rats. Future studies may increase the magnitude of social interaction by increasing the amount of physical contact between rats during the social interaction period. The magnitude of the social reinforcer may also be increased by depriving rats of social interaction during adolescence (Ikemoto and Panksepp, 1992) or adulthood (Chow et al., 2022); however other studies suggest that social depravation during adolescence reduces the reinforcing magnitude of social interaction in adulthood (van den Berg et al., 1999). Future studies should evaluate the degree to which these and other environmental factors affect social reinforcer magnitude in a choice context.

Reduction of the social reinforcer magnitude by keeping the guillotine door closed and removal of the “Partner” rat failed to significantly increase cocaine choice. This result was unexpected, as failure to open the dividing guillotine door and partner rat removal had previously been shown to decrease responding when social access was the only reinforcer available (Baldwin et al., 2022). One potential explanation for the lack of robust effect of removing the social reinforcer is that association between the social stimulus light and social interaction with the “Partner” rat was not sufficiently extinguished over the five experimental days and longer extinction training might be required. Alternatively, stimuli other than the guillotine door opening and presentation of the social rat (e.g., olfactory or visual stimuli) may have contributed to the maintenance of responding on the social-lever. Furthermore, the low level of operant responding for social interaction exhibited in this procedure (about 3 reinforcers/session when 30-s social interaction was available) may have resulted in a floor effect where the current procedure was unable to detect a decrease in social interaction choice. Individual rats did show trends towards either decreased social interaction choice or increased cocaine choice under the 0-s condition, but this did not translate to any significant group differences. Thus, the present results and other studies (Baldwin et al., 2022; Calcagnetti and Schechter, 1992; Chow et al., 2022; Ikemoto and Panksepp, 1992; Trezza et al., 2009; van den Berg et al., 1999) suggest the reinforcing properties of social interaction involve complex interactions with the environment that pose interpretative challenges when designing experiments to manipulate social reinforcer magnitude.

### 4.3 Non-contingent reinforcer presentation did not alter cocaine-vs-social choice

Previous drug-vs-food choice studies have demonstrated that short intertrial intervals enhanced cocaine-vs-saccharin choice in rats (Vandaele et al., 2016). In addition, removal of non-contingent reinforcer presentations before a non-discrete-trial drug-vs-food choice procedure attenuated subsequent drug choice in rats, suggesting that these non-contingent reinforcer presentations functioned as more salient discriminative stimuli than the visual discriminative stimuli presented (Thomsen et al., 2013; Townsend et al., 2021). Together, these findings have been interpreted to suggest that cocaine choice may be enhanced by, or may even require, “choosing under the influence” of drug effects (Vandaele et al., 2016). However, the present results do not support a major role for reinforcer-induced discriminative stimuli and “choosing under the influence” as determinants of cocaine-vs.-social choice in rats. In this study, there was no change in cocaine-vs-social choice when the non-contingent reinforcer presentations before the discrete-choice trials were removed. One potential explanation for this result is the procedural difference between the current discrete-trial cocaine-vs-social choice and prior non-discrete cocaine-vs-food choice studies. In the current discrete-trial procedure, only one cocaine dose was assessed for five experimental days and infusions were separated by 10 min. In previous cocaine-vs-food choice studies by our lab and others (Thomsen et al., 2013; Townsend et al., 2021), multiple cocaine doses were assessed during a single behavioral session and rats could quickly earn up to 10 cocaine infusions within 20-min components. Discriminative stimuli such as lever position and light illumination were possibly more salient in the current discrete-trial cocaine-vs-social choice procedure than in prior non-discrete cocaine-vs-food choice studies. In addition, individual subject behavioral allocation within a daily choice session demonstrated a degree of switching between cocaine and social interaction that appears be related to the unit cocaine dose available rather than any programmed discriminative stimuli.

## 5.0 Conclusions

In summary, the current study found that cocaine availability under a discrete-trial PR schedule was chosen over a social alternative in rats. Cocaine choice was not affected by changes in social-access time as a strategy to manipulate social-reinforcer magnitude, and cocaine choice was not dependent on non-contingent cocaine- and social-reinforcer “sample” presentation preceding choice trials. These findings extend the range of conditions under which cocaine has been shown to maintain choice over an alternative non-drug reinforcer to social interaction in rats. Insofar as drug-choice procedures provide a useful family of tools for preclinical research on the expression, mechanisms, and mitigation of drug reinforcement (Lamb and Ginsburg, 2018; Venniro et al., 2020) the cocaine-vs.-social choice procedure in the present study may have utility in addressing these research questions. Future studies determining the sensitivity of cocaine-vs-social choice to pharmacological treatments such as amphetamine maintenance will be vital to further validate the utility of this procedure in preclinical cocaine addiction research.

## Acknowledgements and Role of the funding source

Research reported in this publication was supported by the National Institute on Drug Abuse of the National Institutes of Health under Award Numbers T32DA007027 and P30DA033934. The content is solely the responsibility of the authors and does not necessarily represent the official views of the National Institutes of Health. Research was also supported by a Virginia Higher Education Equipment Trust Fund Award. Neither funding source had any role in the experimental design, interpretation, or decision to publish the results.

## CRediT author statement

**Madison Marcus:** Methodology, Software, Formal analysis, Investigation, Data curation, Writing – Review and Editing, Visualization. **Steve Negus:** Writing – Review and Editing, Funding acquisition. **Matthew Banks:** Formal analysis, Writing – Original Draft, Writing – Review and Editing, Visualization, Conceptualization, Supervision, Funding acquisition.

